# Ontogeny and expression profiles of steroid hormone receptors in a mouse model of endometriosis

**DOI:** 10.1101/588392

**Authors:** Anuradha Mishra, Mosami Galvankar, Neha Singh, Shantashri Vaidya, Uddhav Chaudhari, Deepak Modi

## Abstract

Endometriosis is a chronic incurable disorder of unknown etiology affecting a large proportion of women in reproductive age. In order to understand the pathogenesis and preclinical testing of drugs,animal models that recapitulate the key features of the disorder are highly desirous. Herein, we describe the ontogeny of the ectopic endometrial lesion in a mouse model where uterine tissue was ligated to the intestinal mesentery and the animals were followed up from day 5 to day 60 post-surgery. Out of 60 animals that underwent surgery, 58 developed endometriosis using this strategy. Most lesions were pale, fluid filled while red lesions were seen in ~10% of animals. Histologically, in most animals there was one large cystic gland with well differentiated epithelium, in 13% of animals there was mixed phenotype (well and poorly differentiated). There was extensive stromal compaction and increased number of macrophages in ectopic lesions. During the course of endometriosis, there was an increase in number of PCNA positive epithelial and stromal cells. The epithelial cells at all the time point were cytokeratin positive and the stroma was vimentin positive. However, at day 30 and 60, the stromal cells were also cytokeratin positive. The mRNA levels of estrogen receptors *Esr1* and *Gper1* were reduced while those of *Esr2* were elevated as compared to normal endometrium, the levels of progesterone receptor (*Pgr*) were found to be downregulated in ectopic lesions as compared to control. However, these differences were not statistically significant due to high biological variability. Low abundance of *Cyp19a1* transcripts (aromatase gene) were only detected in the ectopic endometrium. Immunohistochemically, the expression of ERα and ERβ was significantly reduced only in stromal cells; the epithelial cell staining was maintained. GPER1 and PR immunoreactivity was significantly low in both epithelial and stromal cells. The immunostaining of all the steroid receptors was highly heterogeneous in the ectopic tissues with some areas of sections had stained intensely while others had negligible staining. We propose that temporal and spatial difference in the expression of steroid hormone receptors during the course of endometriosis development coupled with micro-heterogeneity may alter the effectiveness of steroid hormone analogues resulting in variable outcomes and often failure of therapy.

## INTRODUCTUION

Endometriosis is a chronic benign gynecological disorder that results from growth of endometrial glands and stroma outside the uterine cavity. Globally~16.8% of women are estimated to suffer from endometriosis **(Bernuit et al. 2011)**. The symptoms of endometriosis include chronic pelvic pain, dysmenorrhea, dyspareunia and peritoneal inflammation. Laparoscopy is the gold standard method for the diagnosis of endometriosis **(Hsu et al. 2011)** and based on laparoscopic observation the American Fertility Society classified endometriosis into four stages: minimal, mild, moderate and severe stage (**Birmingham et al. 1997)**.

Despite being a common gynecological disorder, the etiology and pathogenesis of endometriosis is still not clear. Effective medical therapies for treatment of endometriosis are lacking (**Tosti et al. 2016)**. Amongst the various possible therapeutic targets, the receptors of the ovarian steroids estrogen and progesterone have taken a central stage. This is because; steroid hormones are known to have a key role in maintaining the endometrial physiology and also proposed to be involved in pathogenesis of endometriosis **(Burney et al. 2012, Barbosa et al. 2011)**. In general, endometriosis is considered to be phenomenon of progesterone resistance and its proliferation is attributed to effect of estrogen **(Bulun et al. 2010)**. Studies have shown that endometriosis is associated with increased local synthesis of estrogen while the expression of progesterone is blunted **(Qi et al. 2017, Kitawaki et al. 2002)**. Based on these observations, anti-estrogen, estrogen receptor modulators and progestin have been evaluated for the therapeutic activity in endometriosis **(Tosti et al. 2017, Rocha et al. 2012 and Taylor et al. 2011)**. However, clinical observations demonstrate none to very moderate effect of these steroid hormone receptors modulators (SERMs) on endometriosis. Wholesome studies have shown very promising effects; others have shown no effects in improving symptoms of endometriosis (**Ferrero et al. 2015**). At present, the reasons for such discrepant findings are unclear.

Estrogen and progesterone act in the target tissue via their receptors mainly the estrogen receptors (ERs) and progesterone receptors (PRs). Several investigators have assessed the expression of ER and PR in endometriotic lesions and the observations vary from study to study. Some studies have shown lower expression of estrogen and progesterone receptors in endometriotic tissue (**Li et al. 2016, Jakson et al. 2007, Nisolle et al. 1994, Bergvist et al. 1993)**; others have shown no difference or higher expression of these receptors in the ectopic endometrium **(Zanatta et al. 2015, Pellegrini et al. 2012, Fujishita et al. 1997, Howell et al. 1994)**. Presently, it is not clear whether such contrasting observations are due to disparity in patient characteristics across the studies or there are inherent differences in the level of these receptors in different types of endometriosis. Furthermore, little is known about the changes that may occur in the receptor’s expression profile during the growth of endometriotic lesions. Considering the importance of steroid hormone therapies in endometriosis, there is a growing need directed towards better understanding of the involvement of steroid hormone receptors in the course of endometriosis.

For development of rational therapeutic strategies, it is essential to have a complete understanding of the disease. While the contributions from human studies are of high value, animal models for endometriosis are highly desirous as they allow us to carry out long term evaluations in controlled conditions. Several non-human primate and rodent models have been described for endometriosis **(Simitsidellis et al. 2018**, **King et al. 2016, and Gonzalez et al 2010)**. Non-human primates can develop endometriosis naturally and have been extensively used to investigate the pathophysiology of endometriosis **(Yamanaka et al. 2012, Dehoux et al. 2011, Braundmeier et al. 2009, Hooghe et al. 2009)**. However, the high cost and difficulty in handling of these animals limit them as experimental model. In this regards, rodents are considered to be most desired animal species not just for ease of handling and cost effectiveness but longitudinal studies on larger numbers of animals can be easily performed **(Bruner-Tran et al. 2018, Grummer et al. 2001)**. In the mouse, xenotransplantation of human tissues into immunocompromised mice have been described as possible model for endometriosis **(Greaves et al. 2017, Bilotas et al. 2015)**. However, this model suffers from the drawback of the being immunocompromised and the role of immune system in development and pathogenesis of endometriosis is well established **(Herington et al. 2011, Grummer et al. 2006)**. To circumvent this, autologous injections of the endometrial tissue intraperitoneally or subcutaneously has beendemonstrated to induce endometriosis like features **(Pereira et al. 2015, Pelch et al. 2012, Grummer et al. 2001)**. However, the lesions developed do not completely resemble the morphological features of endometriosis and the lesions sustain only for a very limited period of time **(Galvankar et al. 2017)**.

At present, surgicalimplantation of autologous endometrial tissue fragment into the peritoneum is by far the most preferred method of inducing endometriosis in rodents. In this model, endometrial tissue fragments are sutured on to the intestinal mesentery which allows growth of the endometrial tissue ectopically. At 30-50 days post-transplant, these lesions have histological and molecularfeatures resembling human endometriosis **(Pereira et al. 2015, Pelch et al. 2012)**. This model has been used for investigating the effect of endometriosis on pain behavior, neuro-visceral interaction, immunological defects and fertility **(Greaves et al. 2017, Bilotas et al. 2015, Somigliana et al. 1999)**.

Despite these applications, the surgical model of endometriosis in the mouse has not been well characterized. In most of the studies, the phenotypes of the ectopic endometrial lesions are studied at a single time point (generally at 30-45 days after induction of the lesions); limited studies have evaluated the evolution and the characteristics of the lesion in animals **(Zhao et al. 2014, Eggermont et al.2005)**. Furthermore, despite being the strong targets for therapy, limited information exists on the spatial and temporal changes in the expression of steroid hormone receptors in the endometrioticlesions inthis model.

In this study, we aimed to study theevolution ofectopic endometrial lesion in a surgically induced autotransplantation mouse model. In addition, we present the spatio-temporal changes in the expression of estrogen receptors (ERα, ERβ & GPER1) and progesterone receptors (PR) in the ectopic endometrium of the mouse model of induced endometriosis.

## RESULS

In all, surgery was performed in 60animals, out of which ectopic lesions were observed in 58 (regressed in two animals) animals resulting in 96.5% success rate.

### Anatomy of the ectopic lesions

Endometrial fragments were surgically implanted on intestinal mesentery and thelesionswereevaluated on day5, 10, 15, 30, 45 and 60 post-surgery (Fig.1A). Macroscopically, ectopic lesionsonday 5, 10 & 15 post-surgery were small but appeared distinct on the intestinal mesentery. The lesions grew in size and by day 30, they appeared large and fluid filled. In most of the animals, lesions were pale and translucent, while in some animals, there werefew hemorrhagic sites within the lesions giving them a reddish appearance (Fig.1B). On day 45 and day 60, the lesion size and appearance remained consistent like that of day 30. Two animals, were studied for120 dayspost-surgery and there were no major changes in its appearance (not shown)

In all the animals, where a single fragment was sutured, one lesion was recovered. In animals, where two fragments were sutured, two lesions were recovered. There was no evidence of de novo lesion development in any of the animals.

Adhesions were not detected in any of the animals on days 5-15 after induction of endometriosis. On day 30, in50% of animals, the lesions had adhesions around the tissue. On day 45 and day 60, all the animals developed dense adhesions attached to the intestine and the lesions were completely buried in them (Fig.1B). At all the time points, the adhesions were always pale white with occasional hemorrhagic sites.

**Figure 1:**
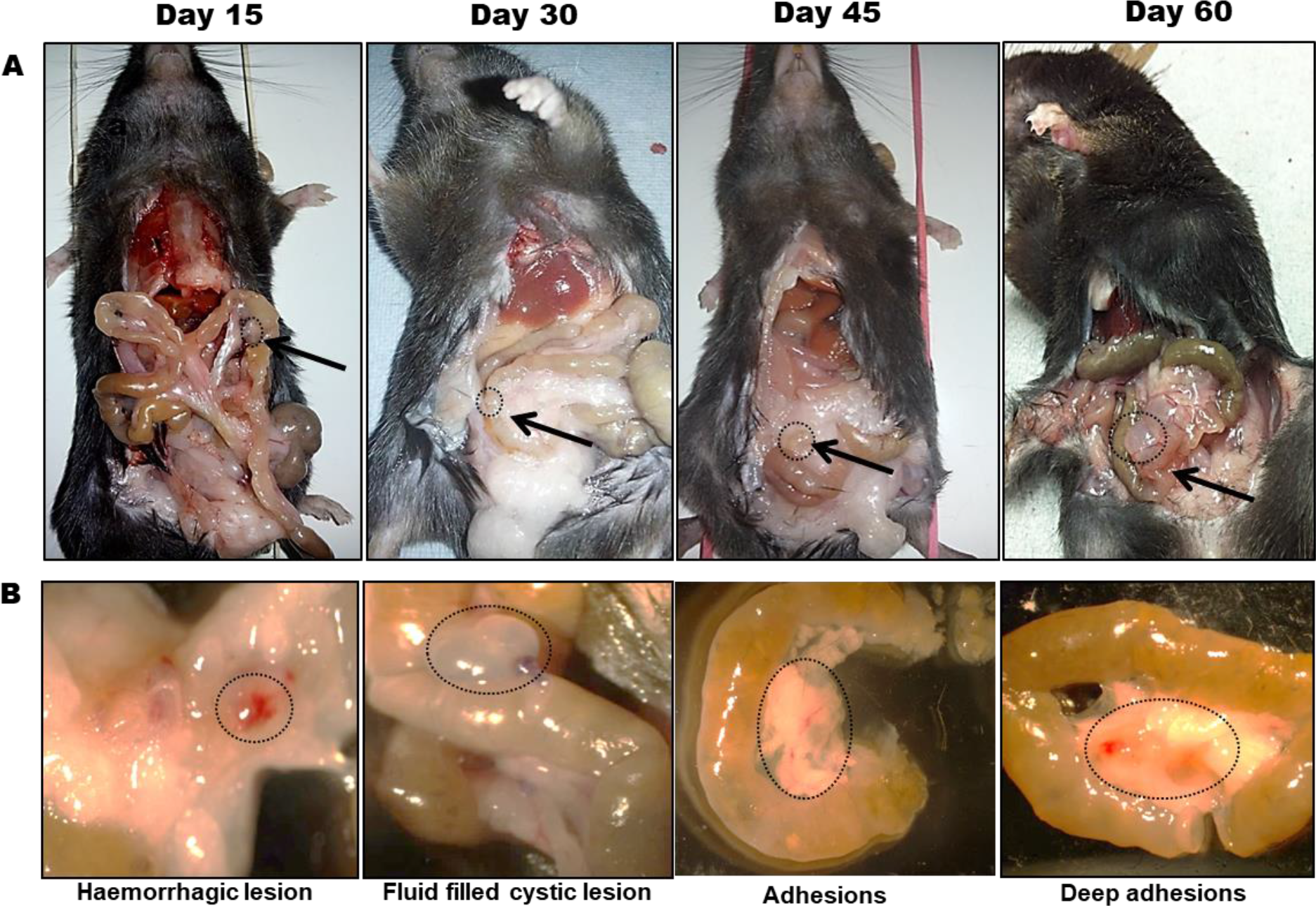
Assessment of endometriotic lesion in surgically induced mouse model. Endometriosis was induced by suturing small uterine fragment to the intestinal mesentery of the same mouse. **(A)** representative unages for lesions developed after 15, 30, 45 and 60 days post-surgery. Circle and arrow showmg lesions developed post-surgery. **(B)** Images showing hemorrhagic & fluid filled lesions with adhesions.

### Histological features of the ectopic lesions

Histologically, theectopic endometrial tissue hadwell-developed glands and stroma at all time points. On day 5, the tissue showed glands and stroma that appeared identical to the control endometrium(Fig.2A). Day 10 onwards, in all the animals, there was a singlecystic gland with large lumen that appeared fluid filled. This cystic gland enlarged in size over time. The epithelial lining sometimes flattened in some areas whereas it was columnar in other areas of the same gland. In tissues collected on day 30 and 45, the cystic glands showed irregular finger like projections similar to lamellae (Fig.2A). By day 60, the cystic glands showed multiple lamellae likeprojections. Epithelium layer appeared thickened and was multilayered in some areas of endometriotic tissue whereas the other glands were small and identical to the endometrium of control animals (Fig.2A).

As compared to control endometrium, in all the ectopic lesions from day 15 onwards thestroma was compactand was hyperplastic. There was no evidence of fibrosis at any time point. In most animals, hemosiderin loaded macrophages were observed in the stroma of ectopic lesions from day10 and further (Fig.2A). No other signs of inflammation were evident in any of the lesions at any time point. Histologically, most of the lesions resembled well-differentiated type of endometriosis; a small subset of animals had mixed appearance with well-differentiated and poorly differentiated lesions; stromal type of histology was never observed in this model (Fig.2B).

**Figure 2:**
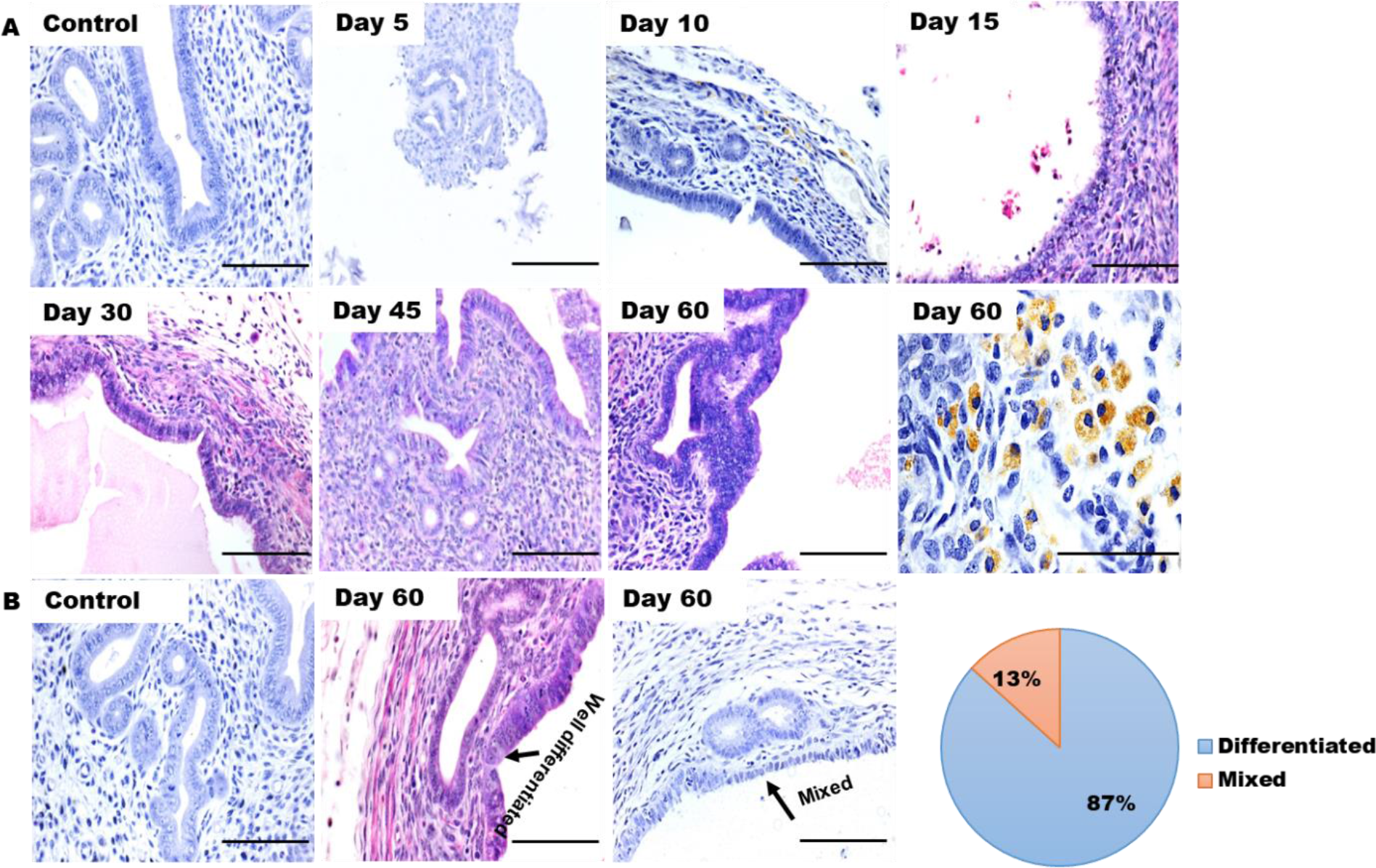
Time dependent changes in the histology of the ectopic lesions in mouse model of endometriosis. Panel A showing representative images of Haematoxylin and Eosin stained paraffin sections of the ectopic lesions on day 5, 10, 15, 30, 45 & 60 post-surgery. Control is endometrium of mice in diestrus stage. Last unage of higher magnification showing hemosiderin laden macrophages in ectopic lesion of day 60 (brown colour). Panel B showing histological pattern of ectopic lesions and pie chart showing percentage of well-differentiated and mixed pattern of endometriosis (n=15); Scale bar, 50μm

### Vimentin and Cytokeratin expression in ectopic lesion

To assess cellular composition of these ectopic lesions, we performed immunohistochemistry for vimentin (stromal marker) and cytokeratin (epithelial marker) in ectopic lesions and in endometrium of control animals. Similar to controls, the stromal cells were strongly stained for vimentin and epithelial cells were stained positive for cytokeratin in the ectopic lesions (Fig.3A & 3C). Quantitatively, as compared to eutopic controls, the expression of vimentin in stroma was marginal but significantly reduced (P<0.05) on day 15. However, on day 30 and 60,expression of vimentin was similar to controls (Fig.3B). As compared to controls, the expression of cytokeratin was significantly (P<0.05) increased in epithelial cells at day 15 and 30. However, by day 60, expression of cytokeratin was similar to endometrium of controls (Fig.3D). At some time points, the stroma appeared to be weak but specifically cytokeratin positive (Fig.3C).

**Figure 3:**
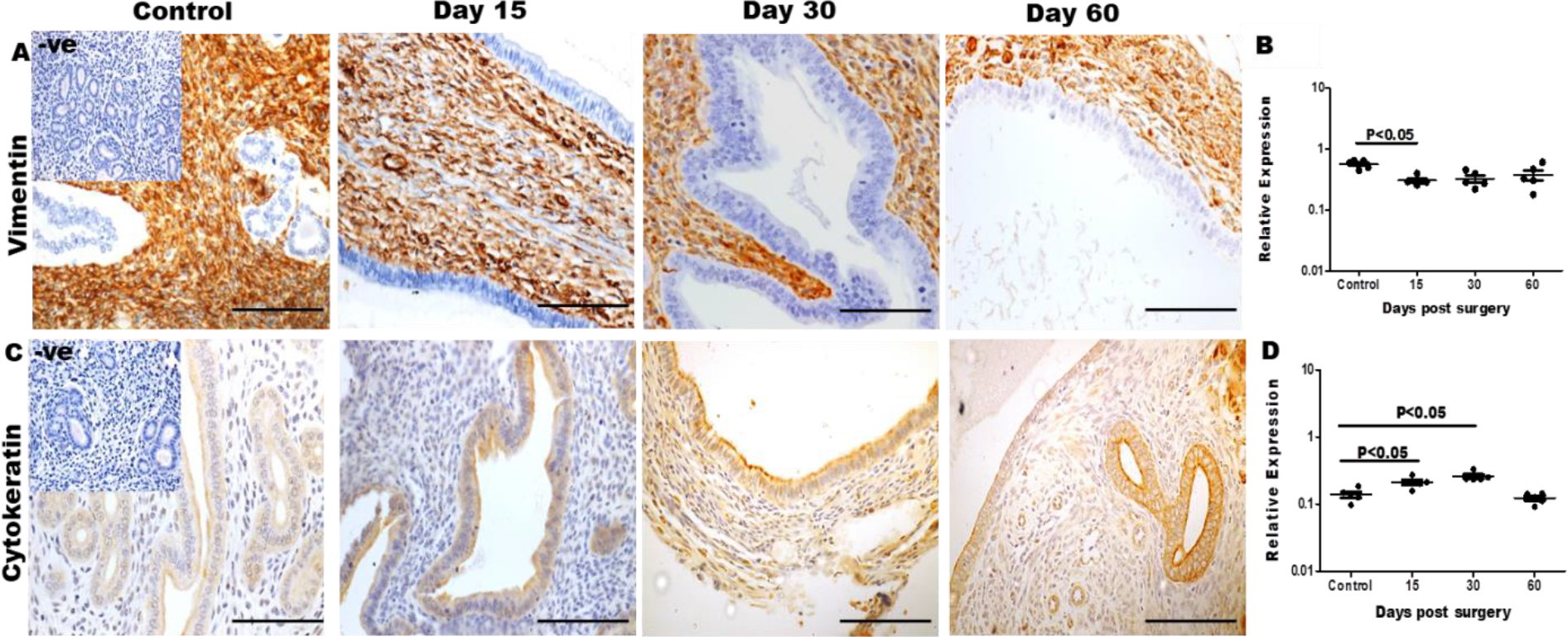
Immunolocalization of vimentin and cytokeratin in ectopic lesions of a mouse model of endometriosis. Ectopic lesions from mice were collected on different days and the tissues sections were stained for vimentin (panel A) and cytokeratin (panel C). Positive staining is indicated by brown stain and blue staining(Haemotoxyllin) is nuclear counterstain. Negative control (-ve) is sections incubated without primary antibody; Scale bar. 5Oμm. Quantification for vimentin (B) and cytokeratin (D) was done and each dot represents intensity value for each animal. Y axis is relative expression of immunostaining in control and ectopic lesions at day 15, 30 and 60. Mean & SD for n=5 control, n=5 ectopic lesion per time point. Statistically significant differences between the groups are shown by horizontal bars.

### Epithelial and stromal cell proliferation is increased in the ectopic lesions

To understand the extent of cell proliferation in the ectopic lesions, we performed immunohistochemistry for PCNA in endometriotic tissues at day 15, 30 and 60 post-surgeryand compared it to controls (Fig.4A). In the control animals at diestrus stage, very few epithelial and stromalcells were positive for PCNA; but in the ectopic lesions, the numbers of PCNA positive cells appeared higher in both glandular epithelium and stroma at all the time points. Quantitatively, there was almost 5-10 fold increase in numbers of PCNA positive cells in epithelium of ectopic lesions as compared to controls at all the time points; this increase was statistically significant (P<0.001). Temporally, the number of PCNA positive cells in the epithelium increased significantly from day 15 to day 60 (Fig.4B).

As compared to controls, there was a significant (P<0.001) increase in PCNA positivestromal cells at all the time points. As compared to day 15, the number of PCNA positive cells was higher in stromal cells on day 30 and 60(Fig.4B).

**Figure 4:**
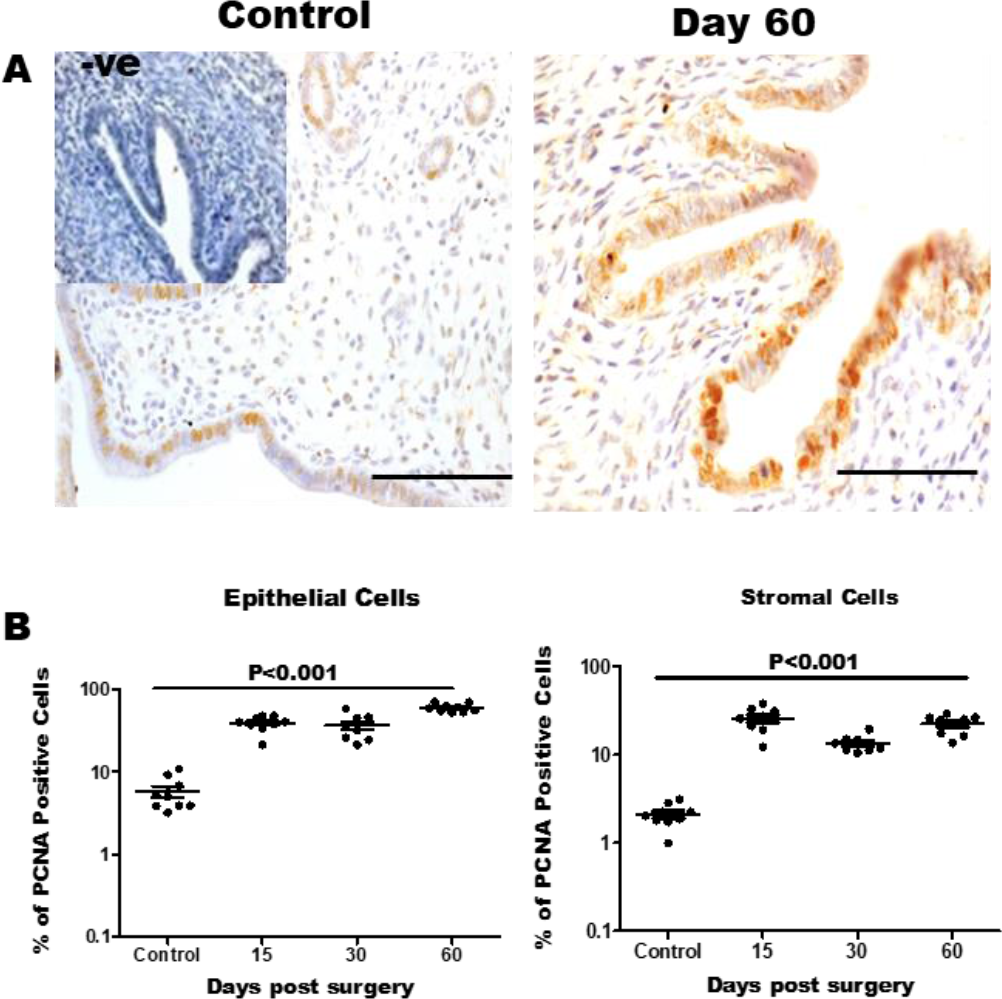
Assessment of proliferation in ectopic endometrial lesions of a mouse model of endometriosis. (A) Immunohistochemistry for PCNA (brown staining) in endometriμm of control mice and ectopic endometrial tissues at day 60 post-surgery. Negative control (−ve) is section incubated without primary antibody (B) Immunostaining quantification for percentage of PCNA positive cell in 3 non-serial sections per animal in both epithelial and stromal cells at day 15, 30 and 60 post-surgery. Mean and SD for n=3 for control, n=3 for ectopic lesion per time point. Scale Bar, 5Oμm. Each dot represents % PCNA positive cells in each area per section per animal. Y axis is % of PCNA positive cells in control and ectopic lesions

### Aromatase gene expression is altered in ectopic endometrium

Abundant *Cyp19a1* mRNA was detected in the ovary (used as positive controls) but not in the control endometrium indicating the specificity of our amplification. However, low abundance *Cyp19a1* transcripts were detected in the endometriotic tissues at all time points. Quantitatively, in the ectopic lesions, *Cyp19a1* transcripts were highest on tissues obtained at day 15, which declined as time progressed. As compared to day 15, the expression of *Cyp19a1* was 1.5folds low on day 30and almost absent on day 60 (Fig.5A).

### Kinetics of mRNA expression of steroid hormone receptors in ectopic endometrial lesion

To assess thechanges in expression of steroid hormone receptors, we performed quantitative real time PCR (qPCR) for steroid hormone receptor genes (*Esr1, Esr2, Gper*and *Pgr)* in diestrus stage ectopic endometrial tissues on day 15, 30 and 60 post-surgery. The eutopicendometriumin diestrus stage was used as control (Fig.5).

#### Estrogen receptorgene1 (*Esr1*) and Estrogen receptor gene2 (*Esr2*)

As compared to controls, ectopic endometrial lesions had 10 foldhigher expression of *Esr1* on day 15which decreased to 5 fold on day 30 as compared to controls. Further, at day 60, mean level of *Esr1* increased to 3 fold as compared to controls. However, this increase was not statistically significant any time point(Fig.5B).

Expression of *Esr2* was increased across all the time points in endometriotic tissues as compared to controls.There was almost 50-foldincrease inmean mRNA levels of *Esr2*at all three days (Day 15, 30 & 60). However, at any of the time point these changes were not statistically significant and there was a high variation across biological replicates (Fig.5C).

Although the mean levels of*Esr1* and *Esr2* levels were higher as compared to control; the levels for both *Esr1* and *Esr2* were higher only in 30-40% animals while the levels were identical or lower than controls in 60-70% animals.Thus, it appears that the failure to achieve statistical significant results for both *Esr1* and *Esr2* isdue to wide variations across the biological replicates.

#### G protein-coupled estrogen receptor gene (*Gper*)

As compared to controls, *Gper* mRNA levels were lower in endometriotic lesionson day 15 and 30 which reached close to normal level on day 60. However, except at day 30, the differences were not statistically significant(Fig.5D). Similar to *Esr1* and *Esr2* mRNA levels, there was variation in the levels of *Gper* across biological replicates. However, these variations were less dramatic as compared to *Esr1* and *Esr2*in endometriotic tissues.

#### Progesterone receptor gene (*Pgr*)

The mean levels of *Pgr* mRNA were identical between endometrium of controls and endometriotic tissues on day 15. *Pgr* mRNA levels were almost 10 fold lower in the endometriotic tissue than controls on day 30 and day 60. Irrespective of these changes, the difference in the mRNA levels between endometriotic tissues and controlswere not statistically significant (Fig.5E).

**Figure 5:**
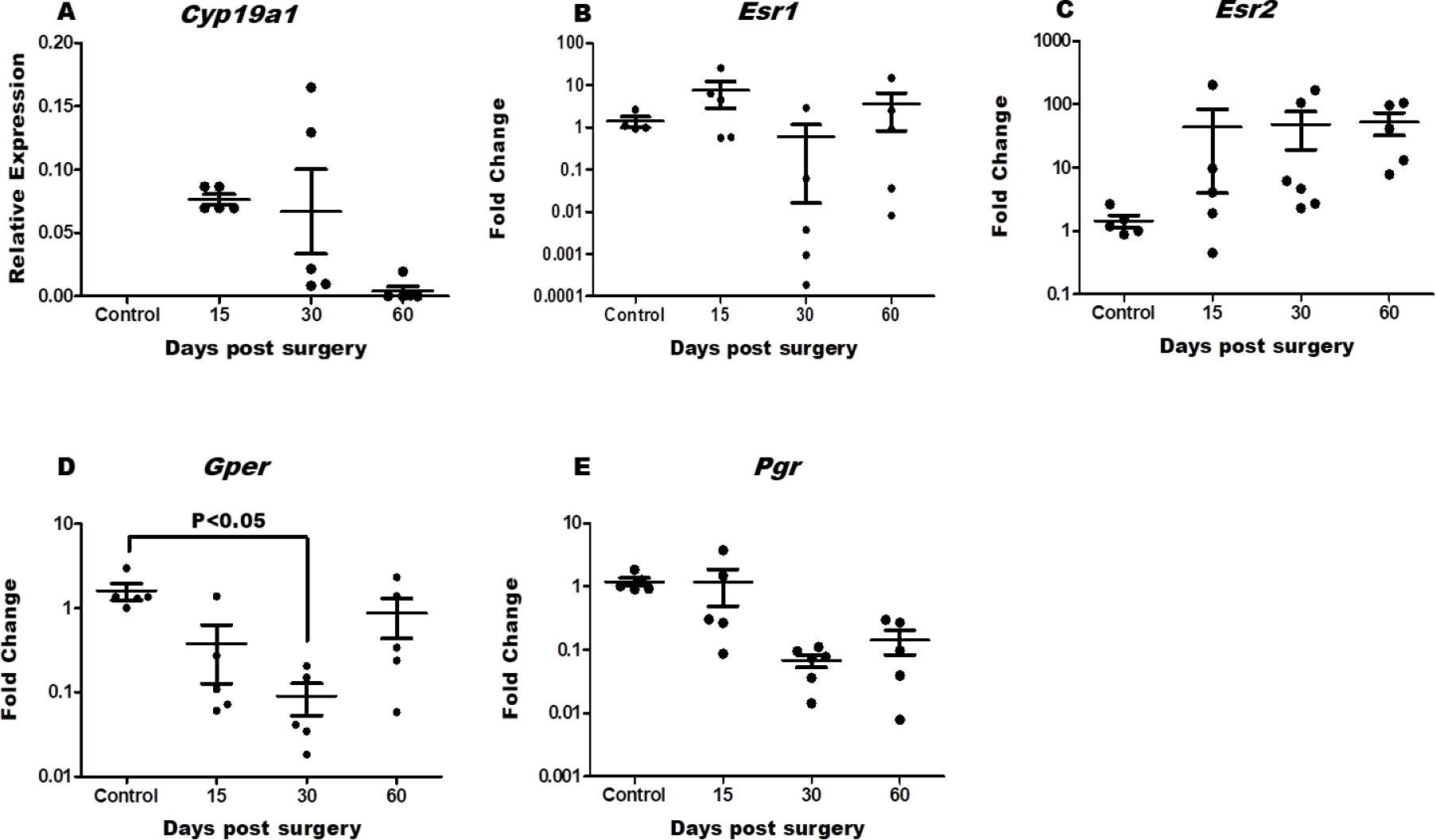
Temporal change in mRNA levels of steroid hormone receptor genes in ectopic endometrial lesions in a mouse model of endometriosis. mRNA levels of *Cup19a1, Esr1, Esr2, Gper* & *Pgr* were quantified by real time PCR in endometrium of control mice and — ectopic endometrial tissues at day 15, 30 & 60 post-surgery. Values on Y axis in *Cup19a1* are relative expression of gene in control and ectopic lesions In *Esr1, Esr2, Gper* & *Pgr*, values on Y axis are fold change in mRNA levels where mean value of control was taken as 1. Each dot represents one animal; the mean and SD for each group is shown (n=5 for control, n=5 for ectopic lesions at each time point)

### Altered spatio-temporal kinetic in ectopic endometrium

Immunohistochemistry was performed to study localization and abundance of ERα, ERβ, GPER1 and PR proteins in endometriotic tissues on day 15, 30 and 60 post-surgery at diestrus stage. Endometrial tissue in diestrus stage was used as control.

#### Estrogen receptor alpha (ERα)

In the endometrium of control animals, the expression of ERα was generally nuclear and appeared higher in epithelial cells as compared to stromal cells. As compared to controls, in endometriotic tissues, epithelial expression of ERα appears reduced on all days while stromal expression of ERα was maintained (Fig.6A).

Quantitative estimation revealed that as compared to controls, the ERα levels were marginally lower on day 15, 30 and 60; however, this reduction was not statistically significant (Fig.7A). To test if ERα is differentially regulated in the epithelial versus stromal cells, quantifications were done separately in both the cell types. In the epithelial cells, as compared to controls, the expression of ERα in the endometriotic tissue was reduced at day 15 and was identical to controls at day 30 and 60. In the stromal cells, the expression of ERα was reduced at all time points as compared to control. However, this decrease was statistically significant (P<0.05) only at day 15 as compared to control (Fig.7A).

#### Estrogen receptor beta (ERβ)

In the endometrium of control animals, ERβ was detected in the cytoplasm of the epithelial cells; the expression in stroma was negligible. In the endometriotic tissues, ERβ often appeared nuclear in both stromal and epithelial cells; the intensity of ERβ staining was also higher in ectopic endometrium as compared to controls (Fig.6B).

As compared to controls, the expression of ERβ was almost 2-5 fold higher in the endometriotic tissues on day 30 and 60. However, this increase was not statistically significant (Fig.7B). In epithelial cells, the expression of ERβ was marginally lower than controls at all the time points. In the stromal cells, as compared to controls, levels of ERβ in ectopic endometrium were higher by 1.5-3 folds on all days (day 15, 30 & 60). However, this increase was statistically significant (P<0.05) only at day 15 as compared to controls (Fig.7B).

#### G protein-coupled estrogen receptor (GPER1)

In endometrium of control animals, GPER1 staining was strong in the epithelium and weak in the stroma. The expression was cytoplasmic and appeared higher on the apical membrane of epithelial cells. As compared to eutopic endometrium of control animals, the intensity of GPER1 appeared lower in the endometriotic tissues (Fig.6C). Quantitatively, ~5 fold reduction in the levels of GPER1 in the endometriotic tissues as compared to controls on all days (15, 30& 60). This reduction was statistically significant (P<0.05) at all the time points (Fig.7C).As observed in Fig.7C, GPER1 protein was significantly (P<0.05) lower in both epithelial and stromal compartment of the endometriotic tissue as compared to control. In both cell types, the reduction of GPER1 was ~ 5-10 fold and statistically significant (P<0.05) on all three days.

#### Progesterone receptor (PR)

PR was detected in the nucleus of both epithelial and stromal cells of the eutopic endometrium. In the ectopic endometrium, the intensity of PR staining was low both in the epithelial and stromal cells, in most cases, the staining was cytoplasmic. As compared to controls, expression of PR was significantly (P<0.05) lower in the endometriotic tissues at all the time points. This reduction was almost 5-10 folds. In almost 50% of animals, there was extremely low PR expression in the endometriotic tissue as compared to controls (Fig.6D & 7D). Levels of PR were also quantified separately in epithelial and stromal cells. As compared to controls, expression of PR was lower in epithelial cells of ectopic endometrium on day 15 and 30 which declined further on day 60. When compared to controls, this reduction was statistically significant (P<0.05) at all the time points. In case of stromal cells, the expression of PR was lower as compared to controls. Quantitatively, there was almost 5-10 fold reduction in the expression of PR. This reduction of PR in the stromal cells of ectopic endometrium was statistically significant (P<0.05) as compared to controls. The downregulation of PR was more dramatic in the stromal cells as compared to epithelial cells in endometriotic tissue.

**Figure 6:**
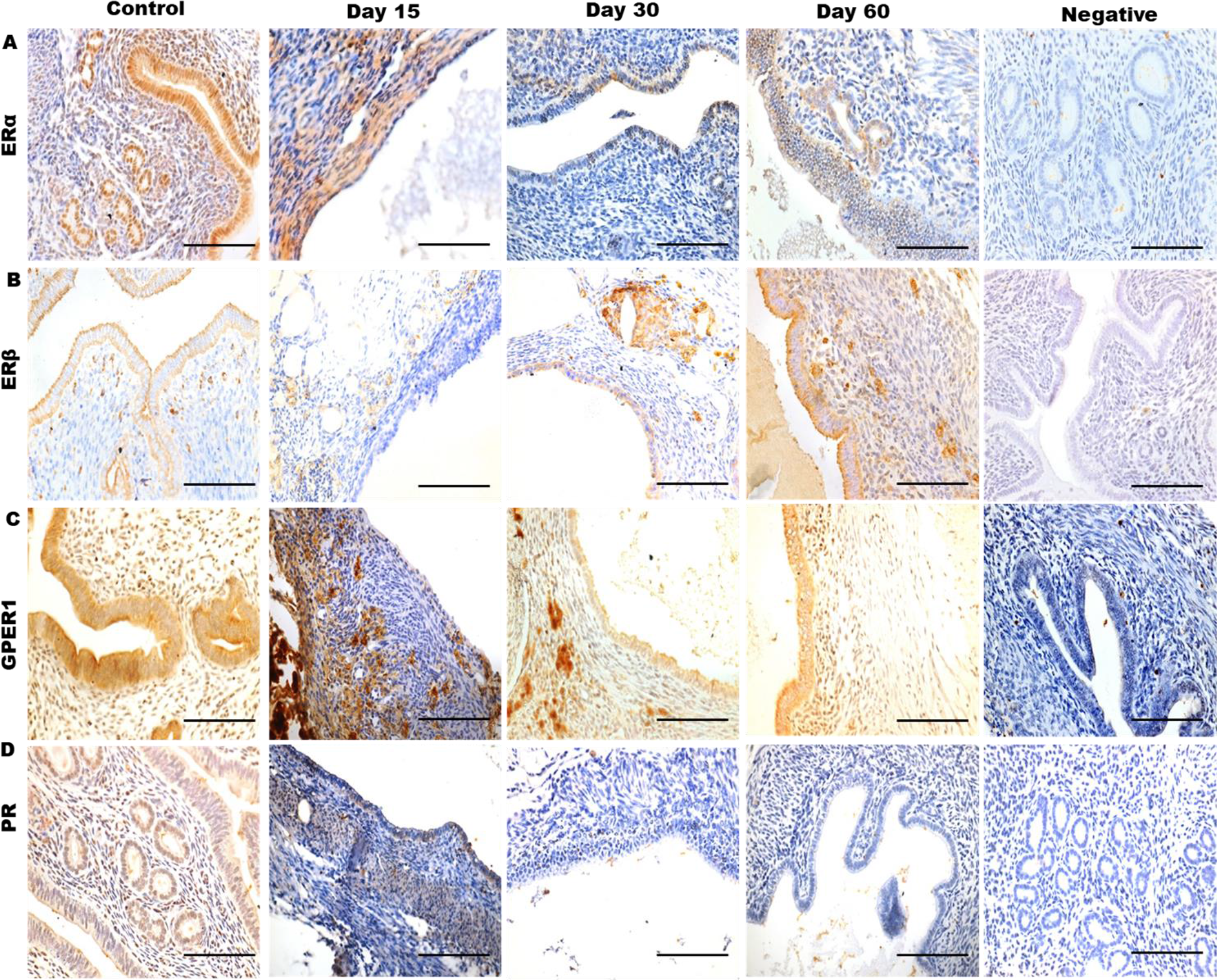
Immuno-localization of steroid hormone receptors in ectopic endometrial lesions of a mouse model of endometriosis. (A-D)Lumunohistochemistry for ERα, ERβ, GPER1 & PR was perfonned in endometriuin of control mice and ectopic endometrial lesions at day 15, 30, 60 post-surgery. Brown staining is indicative of a positive reaction. Cells are counterstained with haematoxylin (blue staining). Scale bar is equivalent to 50 tm. Negative control is section incubated without primary antibody

**Figure 7:**
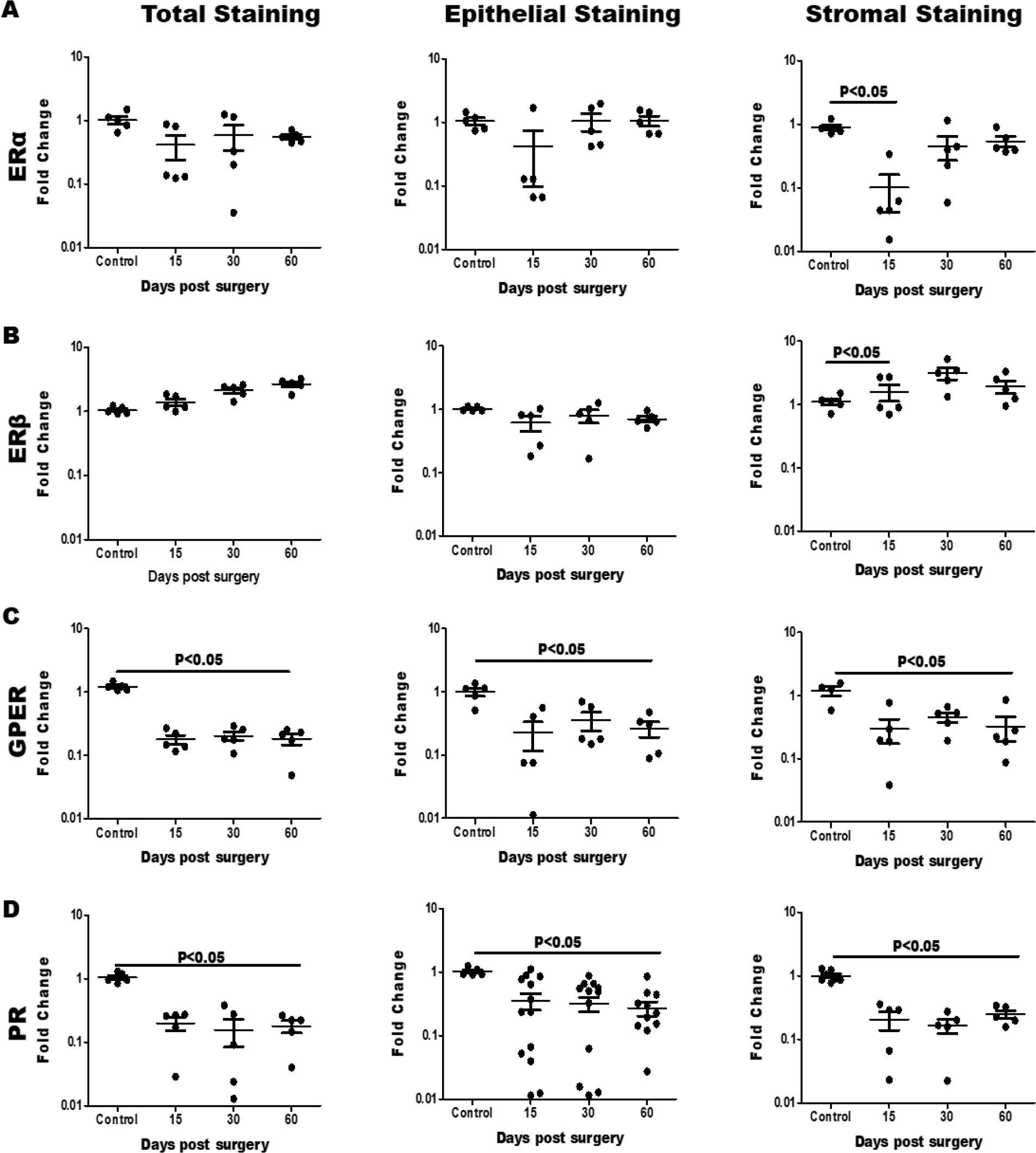
Quantitative changes in levels of steroid hormone receptors in epithelial and stromal cells of ectopic endometrial lesions in a mouse model of endometriosis. (A-D) Immunostaining quantification for ERα, ERβ, GPER1 & PR in endometrium of control mice and ectopic endometrial lesions in whole section and separately in epithelial and stromal cells at 15, 30 and 60 days post-surgery. Each dot represents intensity value for each animal. Values on Y axis are fold change, mean value of control was taken as 1. Mean and SD for n=5 control, n=5 ectopic lesion at day 15, 30 and 60 post-surgery. Statistically significant differences between the groups are shown by horizontal bars.

#### Micro-heterogeneity in expression of steroid hormone receptors in ectopic endometrial lesion

From the above result, we observed that the expression of steroid hormone receptors although altered in the endometriotic tissues as compared to controls; at many instances the differences were not statistically significant and there was a high variability across biological replicates. (Fig.7). During the quantification of immunostaining of steroid hormone receptors in ectopic lesions, we observed that intensity of staining was not uniform across the entire tissue section of lesion while uniform staining was observed in the endometrium of control animals (Fig.8). Also, the expression of cytokeratin and vimentin was found to be uniform across all the sections from ectopic location with little variations between the animals. Thus, it is logical to assume that these changes are not due to technical differences in tissue processing (Fig.3). Hence, we asked if the high biological variability observed in the endometriotic tissue is due to the intra-lesion variability.

To address this question, we quantified the intensity of immunostaining for the steroid hormone receptors in 10 random areas of 3 non-serial sections of endometriotic tissue from three biological replicates onday 60post-surgery andcompared the intensity values in each area. As observed in Fig.8, in the controls, the expression of ERα, ERβ, GPER1 and PR were more or less uniform across the tissues and there was little inter sample variability. However, there was high variability in intensity of staining in different areas of same section of endometriotic tissue and this was observed across all biological replicates. For both ERα and ERβ, the expression in some areas of the endometriotic tissue sectionswas more than the controls; in other areas of the same sections the expression was lower than controls (Fig8A & 8B). Similarly, in case of GPER1 &PR, there was high heterogeneity in the intensity of the staining across the sections obtained from ectopic tissue; the expression was; however, always lower than control animals (Fig.8C & 8D).

**Figure 8:**
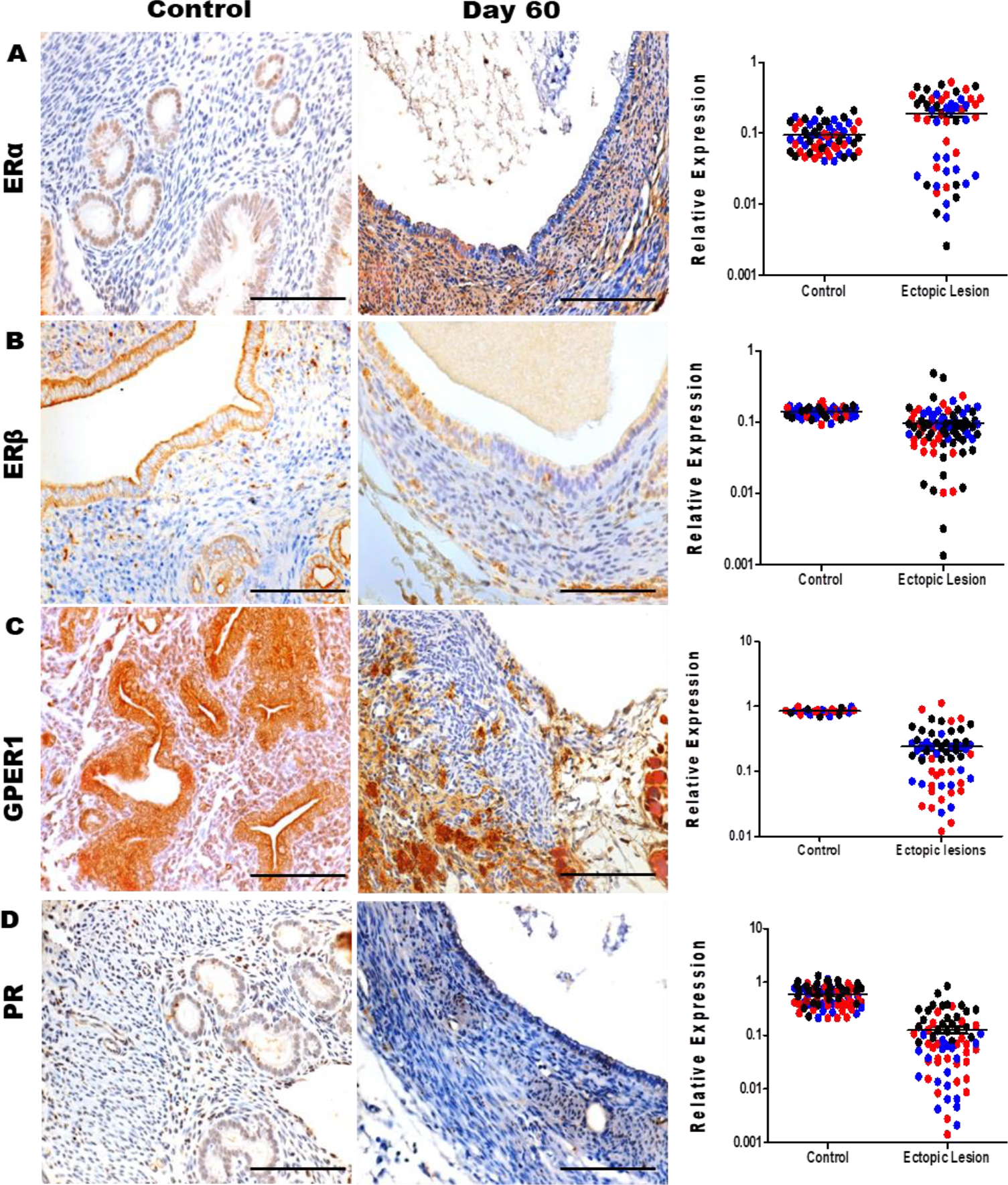
Micro-heterogeneity in expression of steroid hormone receptors in the ectopic endometrial lesion of a mouse model of endometriosis. (A-D) Immunohistochemistry and Immunostaining quantification of ERα, ERβ, GPER1 and PR wm endometrium of control mice and ectopic endometrial tissue at day 60 post-surgery. Brown staining is indicative of a positive reaction. Cells are counterstained with haematoxylin (blue staining). Scale bar, 50 μm. Each dot in graph represents intensity value of unmunostaiming for 10 random areas of 3 non-serial sections per animal. Y axis is relative expression of immunostaining in control and ectopic lesions at day 60. Different colours are given to control and ectopic lesion to show spread of dots from mean value. Mean & SD for n=3 control and n=3 for ectopic lesion. Colour for endometrium of control mice and ectopic endometzial lesions are in blue, red and black.

#### Tissue autonomous regulation of steroid hormone receptors in ectopic endometrial lesion

Clinically endometriosis is characterized by the presence of several lesions in the pelvic peritoneum. We asked if the changes in the expression of steroid hormone receptors observed in endometriotic tissue is lesion independent or the regulation is identical. To test this, two pieces of endometrial tissue were ligated separately onto intestinal mesentery and tissues were excised after 60 days. Both the tissues were independently processed either for qPCR or immunohistochemistry. All the comparisons were made only in the paired tissue of each animal at both gene and protein level.

#### Estrogen receptor genes *(Esr1,Esr2*and *Gper*)

In 6 out of 7 animals, the expression of *Esr1* was discordant between the two lesions obtained from the same animal; in one animal theexpression of *Esr1* between two lesions was identical (Fig.9A). In most of the animals, difference in the expression of *Esr1* between two lesions was in the range of 5-10 folds. The expression of *Esr2*was discordant between two lesions in all the seven animals; in 3/7 animals, there was more than 10 fold differences (Fig.9B).

Similarly, the expression of *Gper* was discordant between the two lesions of same animal. In6/7 animals, the difference in the level of *Gper* between two lesions of same animal was more than 5 fold while in one animal; the level of *Gper* was identical between two lesions (Fig.9C).

#### Progesterone receptor gene(*Pgr*)

In 5/7 animals, the expression of *Pgr* was discordant between two lesions from the same animal; in two animals the levels of *Pgr* mRNA were identical. Amongst the 5 animals that had discordant levels of *Pgr* between the two lesions, in 3 animals the difference was almost10 fold while in other two animals, the difference was more than100 fold (Fig.9D).

Next, we tested if the discordancy in the mRNA levels of steroid hormone receptors between the two lesions also reflected at the protein levels. We compared intensity of ERα, ERβ, GPER1 and PR proteins in two immunostained endometriotic tissue sections of same animal. Like the mRNA, protein levels of steroid hormone receptors were also found to be discordant between the two lesions obtained from the same animal. In general, variation was in the range of 1.5-3 folds between the two ectopic endometrial lesions of same animal (Fig.10).

**Figure 9:**
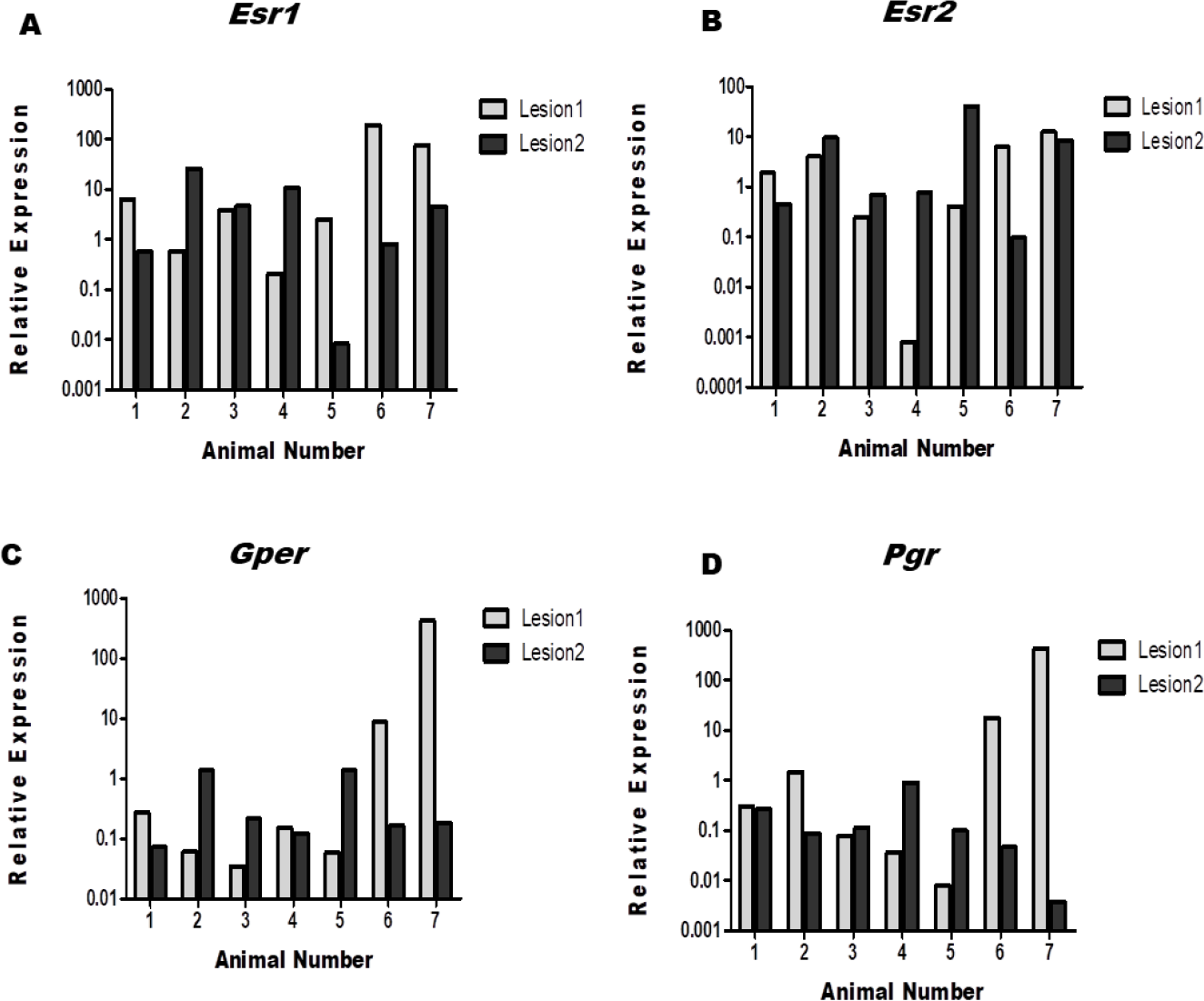
Tissue autonomous regulation of steroid hormone receptor expression in ectopic endometrial lesion of a mouse model of endometriosis. Two endometrial tissue fragments were independently ligated to intestinal mesentery juxtaposed to each other. The lesions were collected on day 60 and independently processed. (A-D) Expression of steroid hormone receptor genes *Esr1, Esr2, Gper* & *Pgr* in two ectopic endometal lesions of the same mouse at day 60 post-surgery by quantitative real time PCR (n=7). Y axis is relative expression of gene in ectopic lesions at day 60.

**Figure 10:**
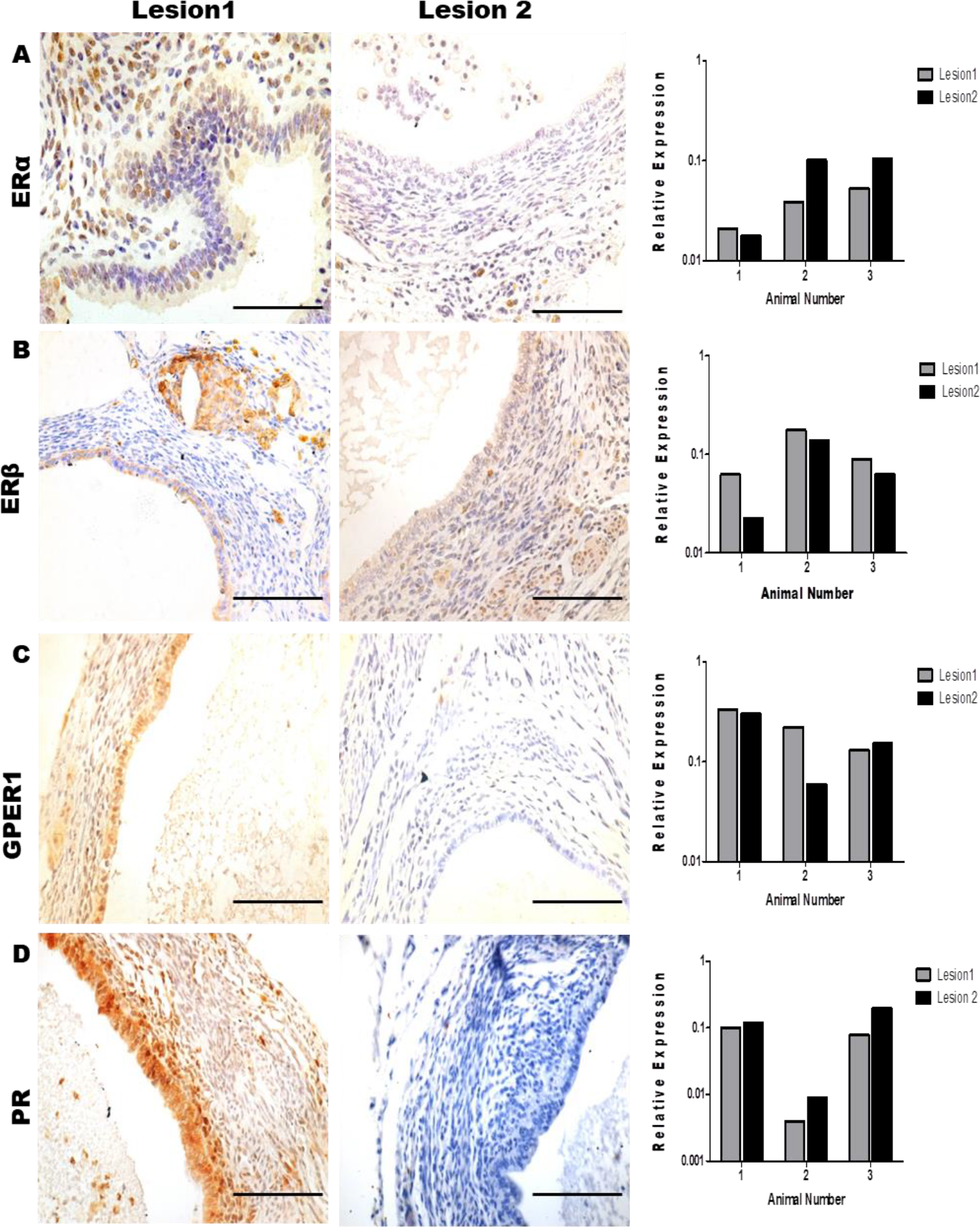
Expression of steroid hormone receptors in ectopic endometrial lesion at protein level. (A-D) Immunohistochemistry and Immunostaining quantification of steroid honnone receptor proteins ERα, ERβ. GPERI & PR in two ectopic endometrial lesions of the same mouse at day 60 post-surgery (n=3). Scale bar, 50μm. Brown staming is indicative of a positive reaction. Cells are counterstamed with haematoxylin (blue staining). Y axis is relative expression of hmmunostaining in ectopic lesions at day 60. mean and SD value for n=3 ectopic lesions at day 60 post-surgery

## DISCUSSION

In the present study, we demonstrate that surgical implantation of the mouse uterine fragments at ectopic location (intestinal mesentery) lead to development of ectopic endometrial lesions. These ectopic lesions grow and progressively acquires histomorphological characteristics similar to human endometriosis. We further show that there is dysregulation in the expression of steroid hormone receptors (ERα, ERβ, GPER1 and PR) in different cell types of endometriotic tissue in a time dependent manner. This dysregulation is not uniform and there is extensive micro heterogeneity in the expression of steroid hormone receptors which seem to be regulated by tissue autonomously.

Animal models of endometriosis are invaluable tools not only in the pharmaceutical industries but also useful for understanding the pathophysiology of the diseases. Several species have been used as animal model for endometriosis **(Simitsidellis et al. 2018, Greaves et al. 2017, King et al. 2016, González et al. 2010 and Grummer et al. 2006)**. Amongst these, mouse model for endometriosis is highly desirable due to the ease of handling and the ability to carry out genetic manipulation. We have previously reported that when fragments of endometrial tissue injected into the peritoneum they tend to adhere at ectopic location but do not sustain for more than seven days even after providing estrogen support **(Galvanker et al. 2017)**. Thus, it was presumed that the mouse endometrial tissue may have poor adhesive capacity to result in endometriosis and requires physical immobilization. In accordance to the previous studies **(Pelch et al. 2012)**, in present the study, we report that surgical immobilization of the endometrial fragments into the intestinal mesentery allows growth with high efficiency. We further show that these lesions survive for long time (almost 6 months post-surgery). During this course, the endometriotic tissue grows in size as a single implant and becomes fluid filled as time progresses. These animals also develop adhesions similar to those observed in women and monkeys **(Fazleabas et al. 2002, Giudice et al. 2010)**.

While the implanted tissue grows in size and develop adhesions like that observed in humans, there are many differences. Inhuman disease and baboon model, the endometriotic lesions not only just grow but also seed new lesions and spread along the peritoneum and also form deep infiltrating lesions if left untreated **(Defrere et al. 2005)**. However, in the surgical model, the endometriotic implants are small, superficial and no new lesions developed in any animal during the study period. Further, in the humans and baboons, the lesions are usually blood filled which often become dark as time progresses and white when regressed **(Fazleabas et al. 2002)**. However, in present study, most of the lesions aretranslucent and pale even in the early stages. In less than 10% of the animals, there is some blood in the lesions.

We next characterized histological ontogeny of the surgically induced endometriotic lesions. In the present study, most of the lesions collected had well-differentiated histomorphology where the epithelium was well defined and often appeared multilayered in some areas indicative of high proliferative activity. However, in 40% of samples, there was a mixed phenotype where in some areas of the glands, the epithelial layer appeared flattened. Mixed phenotypes were generally observed in the lesions obtained on day 45 or day 60 suggesting that the undifferentiated endometriosis seen in a subset of women **(Abrao et al. 2003)** possibly evolved from well differentiated lesions. Irrespective of the phenotype, in all the tissue sections the epithelium stained positive for cytokeratin and negative for vimentin indicating that the composition and organization of the epithelium is well preserved ectopically. This is in contrast to that reported in humans where the ectopic lesions have reduced cytokeratin expression and are vimentin positive suggestive of epithelial to mesenchymal transition (EMT) **(Bartley et al. 2014, Matsuzaki et al. 2012)**. Wealso tested the sections using epithelial and mesenchymal markers and observed that the epithelial cells were E-Cad positive and N-Cad staining was not detected (not shown). These observations suggest that unlike the human condition, EMT is not a feature of endometriosis in the mouse model.

In humans, histological fibrosis is feature of endometriosis observed in nearly 25% of women **(Vigano et al. 2017, Arellano et al. 2011)**. However, none of the animal examined in the present study developed fibrosis despite the lesion being present for more than three months at the ectopic location. In general, the stroma of the endometriotic lesions was hyperplastic at all the time points tested and stained positive for vimentin with no major quantitative differences across time. However, on day 30 and 60, weak but specific staining for cytokeratin was observed in the stromal tissue. Cytokeratin positivity of stromal cells has not been reported previously in endometriosis **(Bartley et al. 2014)**, although it is associated with endometrial stromal sarcoma **(Rahimi et al. 2018)**. However, we did not detect any sarcoma like changes in the ectopic endometrium even at later time points (not shown). The functional significance of such cytokeratin positive stromal cells in the ectopic lesions is unknown.These results imply that the stromal phenotype observed in human endometriosis is not fully recapitulated in this model.

Beyond, epithelial and stromal changes, increased infiltration of macrophages and neutrophils is known to occur in human endometriosis **(Herington et al. 2011, Lin et al. 2006)**. We did observe neutrophils and hemosiderin loaded macrophages in stroma of the endometriotic tissues obtained on day 30 onwards suggestive of inflammation, extensive damage and repair of the tissue in ectopic locations. Together our observations indicate that the mouse model of endometriosis progressively acquire some histological features resembling human endometriosis however marked differences are seen in stromal phenotypes and absence of EMT.

A characteristic feature of endometriosis is extensive proliferation of the tissues at ectopic locations **(Park et al. 2009, Wingfield et al. 1995)**. In concordance with the previous studies using the mouse model, in the present study, we observed that the ectopic lesions have higher numbers of PCNA positive cells as compared to eutopic endometrium with the numbers of proliferative cells increasing with time. Increased proliferation of epithelial cells is observed in the ectopic endometrium of the mouse model as early as one day post-transplantation **(Lin et al. 2006)**; herein we show that the proliferation index increases with time. More than 20% of stromal cells and 60% of epithelial cellsare PCNA positive by day 60. These results suggest that the ectopic endometrial lesion progressively gain a higher capacity to proliferate and therefore as the time progress, these endometriotic tissues not only adherebut alsogrow progressively in size at ectopic location.

Estrogen is a well-known regulator of endometrial cell proliferation and mice lacking ERs specifically ERα have compromised proliferation. In case of endometriosis, several studies have shown that while the systemic levels of estrogen are not significantly different in normal and endometriotic women; however, estrogen is essential for induction of endometriosis in a transplantation mouse model **(Kitawaki et al. 2002, Galvanker et al. 2017)**. It has been shown that the ectopic endometrium gains an ability to locally produce estrogen by expression of aromatase gene. Both, transcripts and protein for aromatase reported in ectopic lesions of women with endometriosis, production of aromatase at ectopic location was also observed in baboons with induced endometriosis **(Qi et al. 2017, Attar et al. 2009)**. In the present study, low but specific transcripts of *Cyp19a1* weredetected in endometriosis as early as day 15 suggesting an increased local synthesis of estrogen. Corroborating this data, a previous study has reported expression of aromatase in 66% of endometrial lesions 4 weeks post transplantation and the growth of the lesion could be suppressed by treatment with aromatase inhibitors **(Bilotas et al. 2010)**. However, it must be noted that herein, we observed a reduction in the abundance of *Cyp19a1* transcripts indicating that local estrogen synthesis may be a feature of early endometriosis, the tissue might lose the ability to produce estrogen with time. This is in contrast with the observations in baboons where aromatase transcripts was detected in the endometriotic lesions only after 10 months of induction which respond to treatment with aromatase inhibitors **(Langoi et al. 2013, Fazleabas et al. 2003)**.

Estrogen acts via its receptors mainly ERα and ERβ. In cyclic endometrium, ERβ is constitutively expressed while the level of ERα is altered with the phase of the endometrial cycle in response to estrogen**(Koehler et al. 2005)**. Functionally, ERα is required for endometrial cell proliferation while ERβ is necessary for regulation of immune responses **(Hen et al. 2015)**. In the context of endometriosis, both ERα and ERβ have been thought to be essential for growth and development of endometriosis **(Barbosa et al. 2011, Bulun et al. 2012)**. However, controversies exist regarding expression profiles of estrogen receptors in the endometriosis. Some studies have shown the increased expression of both ERα and ERβ in the ectopic endometrium of women with endometriosis **(Zanatta et al. 2015, Shao et al. 2014, Pellegrini et al. 2012, Fujishita et al. 1997)** while others have shown the reduced expression of both ERα and ERβ **(Shao et al. 2014, Nisolle et al. 1994, Bergqvist et al. 1993)**. Such conflicting result has also been reported in animal models **(Li et al. 2016, Hen et al. 2015, and Jackson et al. 2007)**. Beyond the classical, ERs, estrogen act via the non-canonical membrane receptor, GPER1 **(Qian et al. 2016)**. In the context of endometriosis, there is a single report demonstrating that the expression for GPER1 is increased in ectopic endometrium of women with endometriosis **(Plante et al. 2012)**. To understand if there are any changes in expression of ERs and GPER1 during the course of disease progression in the mouse model of endometriosis, we analyzed their expression profiles in the ectopic endometrium on day 15, 30 and 60 post-surgery. The results reveal that mRNA levels of *Esr1, Esr2*and*Gper* are altered in the ectopic endometrium during the course of lesion development, however there was high inter-animal variability. We suspected that such differences might arise due altered expression of these molecules in a cell type specific manner and these may not be reflected in whole tissue lysate. Indeed, immunostaining of the tissue sections revealed that the expression of ERα, ERβ and GPER1were reduced in the epithelial cells of endometriotic tissue with more dramatic reduction in levels of GPER1 as compared to ERα and ERβ. Interestingly, in the stromal cells, the expression of ERα and GPER1 was reduced as compared to controls at all the time points, while that of ERβ progressively increased. These results imply that there are cell type specific changesin expression of estrogen receptors in ectopic endometrium and both canonical and non-canonical actions of estrogen are affected in endometriosis.

In the cycling endometrium, progesterone acts via its receptors to induce differentiation required to attain receptivity and promote implantation **(Wetendorf et al. 2014)**. In women with endometriosis, although serum levels of progesterone are similar to those of women without the disease, it is well established that endometriotic lesions (ectopic endometrium) do not respond appropriately to progesterone. This is thought to be because oflower levels of PR in endometrium of women with endometriosis**(McKinnon et al. 2018, Patel et al. 2017)**. Corroborating the human data, in the present study, the expression of PR was also reduced in both epithelium and stroma of the endometriotic tissue as compared to endometrium of control animals. Together these observations imply that the canonical andnon-classical ER & PR signaling are hampered in endometriotic tissue.

In the present study, we observed that irrespective of the receptors there was high inter-sample variability. In a recent study, similar high inter-subject and intra-subject variation in the PR levels was reported in human endometriotic lesions **(Flores et al. 2018)**.In addition to high intra-animal variability, quantitative analysis of different regions of same section reveal that the expression of ERα, ERβ, GPER1 and PR were higher in the certain areas of the sections while almost absent in other area of same section. We termed these variations as micro-heterogeneity and quantified it across the biological replicates. The results revealed that the expression of all these proteins varies as much as 100 fold between adjacent areas of the same section and this was also observed in all the endometriotic lesions examined. This micro-heterogeneity was not due to technical errors during tissue processing as expression of other proteins (vimentin and cytokeratin) was homogenous in the same sections with minimal biological variability across the samples.

We next addressed whether this micro-heterogeneity is driven by systemic clue or it is tissue autonomous. To answer this question, we surgically implanted two pieces of endometrial tissue adjacently in the same animal, harvested 60 days later and processed them independently. The results reveal that there was no concordance in the levels of steroid hormone receptors between the two lesions isolated from the same animal. This was also reflected at the protein levels where one lesion has higher expression than normal while others have lower expression than normal. This was true for all ERα, ERβ, GPER1 and PR implying that micro-heterogeneity observed in the endometriosis is tissue autonomous and is not systemically driven. At present, it is difficult to speculate what would be the reason for such micro-heterogeneity. In endometriosis, altered expression of ER and PR are thought to be due to differential levels of the DNA methylation of their cognate genes in the genome **(Xue et al. 2007, Wu et al. 2006)**. We suspect that such methylation changes may not have uniformly occurred throughout the tissue and this may contribute to such differential expression of these receptors. From the results of the present study, we suspect that the discordant results obtained in different studies on expression of steroid hormone receptors could arise due to site and time of tissue sampling. Thus, one needs to be very cautious while interpreting the data from such studies and concept of micro-heterogeneity must be borne in mind while analyzing the data on endometriotic tissue.

In the context of steroid hormone receptors, our results demonstrate that there is high inter and intra lesion variability which could also alter with time. This implies that the sensitivity to exogenous steroid hormones may not be uniform. Several systematic reviews including the Cochran data bases have observed that despite high quality study design, there is inconsistency in the outcomes of steroidal therapies between different studies **(Bozdag et al. 2015)**. It is possible thatthe inconsistency across studies in response to hormone therapies is not due to methodological issues but is inherent to endometriosis. It is plausible that temporal and spatial difference in the expression of steroid hormone receptors coupled with micro-heterogeneity may alter the effectiveness of steroid hormone analogues resulting in variable outcomes and often failure of therapy.

In summary, the results of the present study have shown that it is possible to partially mimic the stage I-IV of well differentiated endometriosis in the mouse model. However, our results show that there are some differences in the evolution of endometriosis in human subjects and in this system. Furthermore, herein, we demonstrate that there is extensive heterogeneity in the expression of classical and non-classical steroid hormone receptors in the endometriotic tissues which cautious us regarding the therapeutic use of steroid hormone modulators in women with endometriosis. We believe that characterizing the mouse model of endometriosis would be animmense developing rationale for diagnosis and aid in management strategy for this common but as of yet untreatable disorder.

## MATERIAL AND METHODS

### Animal Ethics

This study was approved by the Institutional Animal Ethics Committee (IAEC) of the National Institute for Research in Reproductive (NIRRH) and committee for Biosafety.

### Animals

3-4 month old regularly cyclingC57BL/6 strain mice were used for study. Mice housed in environmentally controlled cages, with 12 h day night cycles.

### Endometriosis induction

Endometriosis was surgically induced in the mice as described previously **(Pelch et al. 2012)**. Briefly, mice in diestrus and estrus stage were anesthetized by ketamine and xylazine.Small incision was given on skin and muscleto expose the uterine horns. The left uterine horn was excised, opened medially and cut into 3 small fragments. Onefragment was sutured with the intestinal mesentery; muscles and skin were sutured back andthe animal was revived in a 37°C warm chamber and put back to the cage. In some experiments, two fragments were sutured individually at a distance on the intestinal mesentery.

### Experimental Tissue collection

Animals were sacrificed on day 5, 10, 15, 30, 45 and 60 post-surgery. The tissue attached to intestinal mesentery was excised and stored in Trizol reagent for RNA extraction or fixedin 4% PFA for paraffin embedding and sectioning. In some cases, more than one lesion was sutured in the same animal; both the lesions were excised and processed individually.For controls, uterus from 3-4 month old female mice were alsocollected in Trizol reagent for RNA extraction and in 4% PFA for paraffin embedding and sectioning.

### RNA Extraction &cDNA Synthesis

Total RNA was extracted using Trizol reagent (Invitrogen), treated with DNase (GE Healthcare, Hongkong, China), and reverse transcribed by Clonetech cDNA synthesis kit as described previously **(Godbole et al. 2007).**

### Quantitative Real Time PCR

Specific primers were designed to check the expression of *Esr1, Esr2, Cyp19a1, Gper* and *Pgr*. The data was normalized to the level of *18srRNA*. The primer sequences for the above genes are given in Table1.Real time PCR was done in triplicates as detailed previously **(Godbole et al. 2007, Lahari et al. 2018)**. All amplifications were done using theCycler Real Time PCR System (Bio-Rad) andSybr Green Chemistry (Biorad). For each primer pair, reaction efficiency was estimated by the amplification of serial dilution of mouse uterus cDNA pool over 5 fold range. The amplification condition for each primer were: initial denaturation at 98°C for 2 minutes followed by 40 cycles of denaturation at 98°C for 30s, primer annealing at their respective optimized T_a_ and extension at 72°C for 45s, melt curve 75°C to 95°C. The fluorescence emitted at each cycle was collected for the entire period of 30s during the extension step of each cycle.

The homogeneity of the PCR amplicons was verified by the melt curve method. All PCR was carried out in triplicates. Mean Cq value generated in each experiment using the iCycler software (Bio-Rad) were used to obtain the standard curve,).The relative expression ratios were calculated using Livak method **(Livak et al. 2001)** and statistically analyzed by one way Anova in GraphPad Prism.

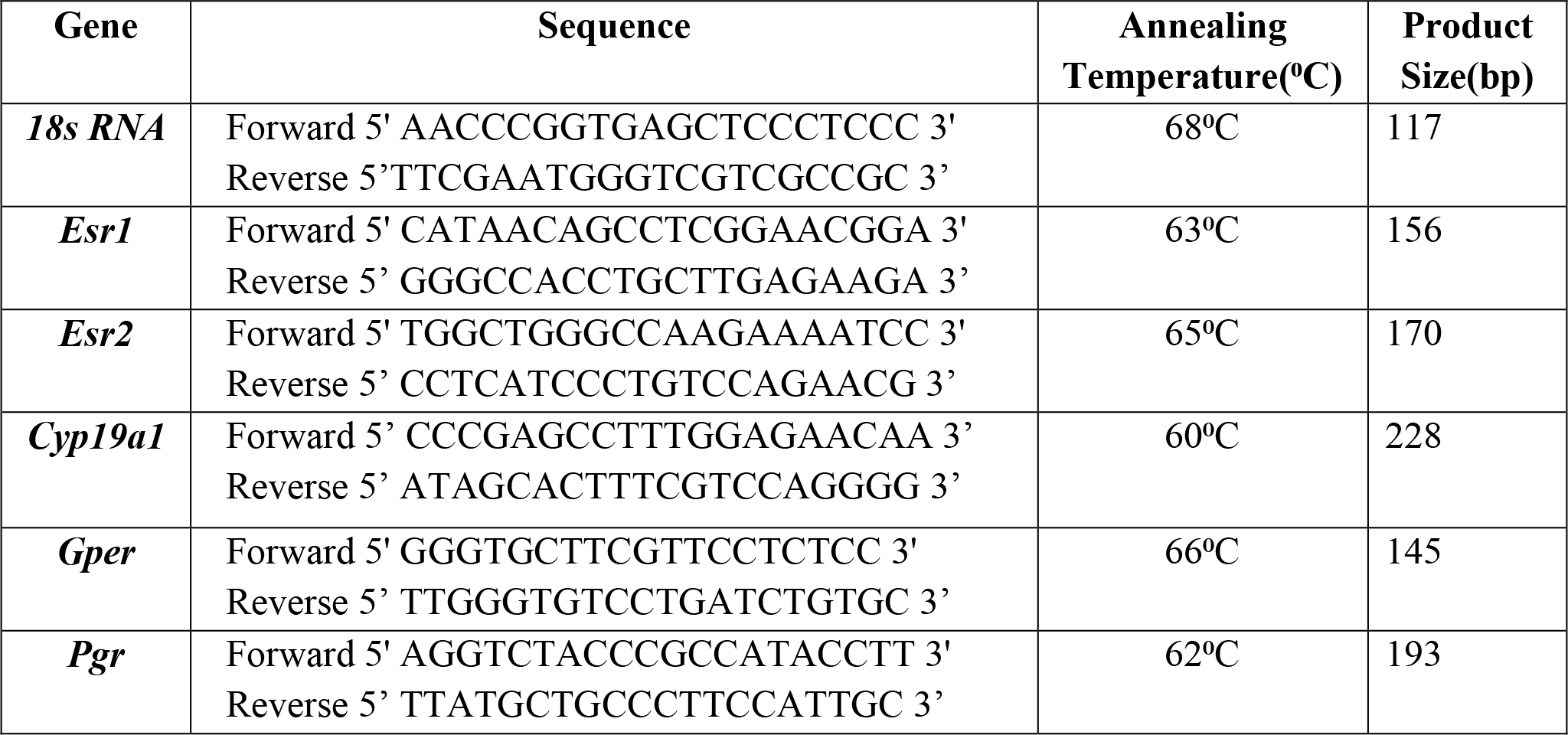

### Hematoxylin & Eosin Staining

Five micrometers thick paraffin sections of 4% PFA fixed tissueswere cut and mounted on poly-L-lysine coated slides. Sections were deparaffinised in xylene, hydrated in descending grades of alcohol for 5 minutes. After that endometriotic tissue sections were stained with hematoxylin stain (Himedia)followed by eosin (Himedia) and mounted. Slides were viewed under bright field microscope (Olympus)and representative areas were photographed.

### Immunohistochemistry

Immunohistochemistry was performed as described previously **(Godbole et al. 2007, Laheri et al. 2017).** Five micrometers thick sections were deparaffinised in xylene, hydrated in descending grades of alcohol. Antigens were retrieved by Tris-EDTA Buffer (pH-9), blocking was done in 5% BSA and the sections were probed overnight with primary antibody. Negative controls were incubated with PBS instead of primary antibody. Next day, slides were washed three times with PBS for 10 minutes and incubated with biotinylated secondary antibody followed by incubation in streptavidin-HRP (ABC Santa Cruz Biotechnology) for 30 minutes. Detection was done using hydrogen peroxidase substrate and Diaminobenzidine (Sigma Aldrich) as chromogen. All sections were briefly counterstained with hematoxylin and mounted. Slides were viewed underbright Field microscope(Olympus) and representative areas were photographed.

### Immunostaining Score

Immunostaining was quantified in terms of intensity numbers by "Fiji" version of ImageJ. To quantify immunostaining for individual proteins in endometrium, we selected three different images showing different area of same lesion and calculated the intensity numbers. For individual cell types, stromal and epithelial cells, we selected randomly 10 different areas of 2000m^2^ in stroma and epithelia and calculated intensity numbers. To quantify microhetergeneity, we selected 10 random areas of 2000m^2^ in 3 non-serial sections of 3 biological replicates and quantified intensity number same as done for stromal and epithelial cells.

This intensity number was further converted into optical density (OD) with the following formula in MS Excel sheet.

Optical Density (OD) = log (max intensity/Mean intensity), where max intensity = 255 for 8-bit images.

### Statistical Analysis

All experimental quantitative data of various genes and proteins was calculated in terms of Mean and Standard deviation, and statistical analysis was done using GraphPad Prism, version 5, either by one-way ANOVA using Tukey’s all column comparison test or Student’s unpaired t test. P < 0.05 was accepted as statistically significant.

## Acknowledgements

We express our gratitude to Dr Stacy Colaco and Mr Abhishek Tiwari for help during preparation of the manuscript. We thank the staff of the Animal House (NIRRH) for their help during the surgeries and animal maintenance.The manuscript bears the NIRRH ID: RA/756/03-2019. DM lab is funded by grants from ICMR, Govt of India. The study was funded by grants from Department of Biotechnology (DBT), Govt of India (BT/OR6587/MED/30/886/2012) to DM. AM is recipient of the junior and senior research fellowship from University Grants Commission (UGC) Govt of India. MG was the recipient of the ICMR postdoctoral fellowship(6^th^Batch).

## References

1. Abrao MS, Neme RM, Carvalho FM, Aldrighi JM, Pinotti J. Histological classification of endometriosis as a predictor of response to treatment. International Journal of Gynecology & Obstetrics. 2003; 82(1):31–40.

2. Attar E, Tokunaga H, Imir G, Yilmaz MB, Redwine D, Putman M, Gurates B, Attar R, Yaegashi N, Hales DB, Bulun SE. Prostaglandin E2 via steroidogenic factor-1 coordinately regulates transcription of steroidogenic genes necessary for estrogen synthesis in endometriosis. The Journal of Clinical Endocrinology & Metabolism. 2009;94; 94(2):623–31.

3. Barbosa CP, De Souza AB, Bianco B, Christofolini DM. The effect of hormones on endometriosis development. Minerva Ginecol. 2011; 63(4):375–86.

4. Barcena de Arellano ML, Gericke J, Reichelt U, Okuducu AF, Ebert AD, Chiantera V, Schneider A, Mechsner S. Immunohistochemical characterization of endometriosis-associated smooth muscle cells in human peritoneal endometriotic lesions. Human reproduction. 2011; 26(10):2721–30.

5. Bartley J, Jülicher A, Hotz B, Mechsner S, Hotz H. Epithelial to mesenchymal transition (EMT) seems to be regulated differently in endometriosis and the endometrium. Archives of gynecology and obstetrics. 2014;289(4):871–81.

6. Bergqvist A, Ljungberg O, Skoog L. Endometriosis: Immunohistochemical analysis of oestrogen and progesterone receptors in endometriotic tissue and endometrium. Human Reproduction. 1993;8; 8(11):1915–22.

7. Bernuit D, Ebert AD, Halis G, Strothmann A, Gerlinger C, Geppert K, Faustmann T. Female perspectives on endometriosis: findings from the uterine bleeding and pain women's research study. Journal of Endometriosis. 2011; 3(2):73–85.

8. Bilotas MA, Olivares CN, Ricci AG, Baston JI, Bengochea TS, Meresman GF, Barañao RI. Interplay between endometriosis and pregnancy in a mouse model. PloS one. 2015; 10(4):e0124900.

9. Birmingham & Alabama. Revised American Society for Reproductive Medicine classification of endometriosis: 1996. Fertility and sterility 1997; 67(5).

10. Bozdag G. Recurrence of endometriosis: risk factors, mechanisms and biomarkers. Women's Health. 2015; 11(5):693–9.

11. Braundmeier AG, Fazleabas AT. The non-human primate model of endometriosis: research and implications for fecundity. Molecular human reproduction. 2009; 15(10):577–86.

12. Bruner-Tran KL, Mokshagundam S, Herington JL, Ding T, Osteen KG. Rodent models of experimental endometriosis: identifying mechanisms of disease and therapeutic targets. Current women's health reviews. 2018;14(2):173–88.

13. Bulun SE, Cheng YH, Pavone ME, Xue Q, Attar E, Trukhacheva E, Tokunaga H, Utsunomiya H, Yin P, Luo X, Lin Z. Estrogen receptor-β, estrogen receptor-α, and progesterone resistance in endometriosis. InSeminars in reproductive medicine 2010:28(01):036–043. © Thieme Medical Publishers.

14. Bulun SE, Monsavais D, Pavone ME, Dyson M, Xue Q, Attar E, Tokunaga H, Su EJ. Role of estrogen receptor-β in endometriosis. InSeminars in reproductive medicine 2012:30(01): 39–45. Thieme Medical Publishers.

15. Burney RO, Giudice LC. Pathogenesis and pathophysiology of endometriosis. Fertility and sterility. 2012; 98(3):511–9.

16. Defrère S, Van Langendonckt A, Ramos RG, Jouret M, Mettlen M, Donnez J. Quantification of endometriotic lesions in a murine model by fluorimetric and morphometric analyses. Human Reproduction. 2005; 21(3):810–7.

17. Dehoux JP, Defrère S, Squifflet J, Donnez O, Polet R, Mestdagt M, Foidart JM, Van Langendonckt A, Donnez J. Is the baboon model appropriate for endometriosis studies?. Fertility and sterility. 2011;96(3):728–33.

18. d'Hooghe TM, Kyama CM, Chai D, Fassbender A, Vodolazkaia A, Bokor A, Mwenda JM. Nonhuman primate models for translational research in endometriosis. Reproductive Sciences. 2009;(2):152–61.

19. Eggermont J, Donnez J, Casanas-Roux F, Scholtes H, Van Langendonckt A. Time course of pelvic endometriotic lesion revascularization in a nude mouse model. Fertility and sterility. 2005;84(2):492–9.

20. Fazleabas AT, Brudney A, Chai D, Langoi D, Bulun SE. Steroid receptor and aromatase expression in baboon endometriotic lesions. Fertility and sterility. 2003;80:820–7.

21. Fazleabas AT, Brudney A, Gurates B, Chai D, Bulun S. A modified baboon model for endometriosis. Annals of the New York Academy of Sciences. 2002; 955(1):308–17.

22. Ferrero S, Alessandri F, Racca A, Maggiore UL. Treatment of pain associated with deep endometriosis: alternatives and evidence. Fertility and sterility. 2015; 104(4):771–92.

23. Flores VA, Vanhie A, Dang T, Taylor HS. Progesterone receptor status predicts response to progestin therapy in endometriosis. The Journal of Clinical Endocrinology & Metabolism. 2018;103(12):4561–8.

24. Fujishita A, Nakane PK, Koji T, Masuzaki H, Chavez RO, Yamabe T, Ishimaru T. Expression of estrogen and progesterone receptors in endometrium and peritoneal endometriosis: an immunohistochemical and in situ hybridization study. Fertility and sterility. 1997; 67(5):856–64.

25. Galvankar M, Singh N, Modi D. Estrogen is essential but not sufficient to induce endometriosis. Journal of biosciences. 2017; 42(2):251–63.

26. Giudice LC. Endometriosis. New England Journal of Medicine. 2010;362(25):2389–98.

27. Godbole G, Suman P, Malik A, Galvankar M, Joshi N, Fazleabas A, Gupta SK, Modi D. Decrease in expression of HOXA10 in the decidua after embryo implantation promotes trophoblast invasion. Endocrinology. 2017;158(8):2618–33.

28. Godbole GB, Modi DN, Puri CP. Regulation of homeobox A10 expression in the primate endometrium by progesterone and embryonic stimuli. Reproduction. 2007;134(3):513–23.

29. Greaves E, Cousins FL, Murray A, Esnal-Zufiaurre A, Fassbender A, Horne AW, Saunders PT. A novel mouse model of endometriosis mimics human phenotype and reveals insights into the inflammatory contribution of shed endometrium. The American journal of pathology. 2014; 184(7):1930–9.

30. Greaves E, Critchley HO, Horne AW, Saunders PT. Relevant human tissue resources and laboratory models for use in endometriosis research. ActaobstetriciaetgynecologicaScandinavica. 2017; 96(6):644–58.

31. Greaves E, Horne AW, Jerina H, Mikolajczak M, Hilferty L, Mitchell R, Fleetwood-Walker SM, Saunders PT. EP 2 receptor antagonism reduces peripheral and central hyperalgesia in a preclinical mouse model of endometriosis. Scientific reports. 2017; 7:44169.

32. Grümmer R, Schwarzer F, Bainczyk K, Hess-Stumpp H, Regidor PA, Schindler AE, Winterhager E. Peritoneal endometriosis: validation of an in-vivo model. Human Reproduction. 2001; 16(8):1736–43.

33. Grümmer R. Animal models in endometriosis research. Human reproduction update. 2006;12(5):641–9.

34. Herington JL, Bruner-Tran KL, Lucas JA, Osteen KG. Immune interactions in endometriosis. Expert review of clinical immunology. 2011;7(5):611–26.

35. Howell RJ, Dowsett M, Edmonds DK. Endometriosis: Oestrogen and progesterone receptors in endometriosis: heterogeneity of different sites. Human Reproduction. 1994; 9(9):1752–8.

36. Hsu AL, Khachikyan I, Stratton P. Invasive and non-invasive methods for the diagnosis of endometriosis. Clinical obstetrics and gynecology. 2010; 53(2):413.

37. Jackson KS, Brudney A, Hastings JM, Mavrogianis PA, Kim JJ, Fazleabas AT. The altered distribution of the steroid hormone receptors and the chaperone immunophilin FKBP52 in a baboon model of endometriosis is associated with progesterone resistance during the window of uterine receptivity. Reproductive Sciences. 2007 (2):137–50.

38. King CM, Barbara C, Prentice A, Brenton JD, Charnock-Jones DS. Models of endometriosis and their utility in studying progression to ovarian clear cell carcinoma. The Journal of pathology. 2016; 238(2):185–96.

39. Kitawaki J, Kado N, Ishihara H, Koshiba H, Kitaoka Y, Honjo H. Endometriosis: the pathophysiology as an estrogen-dependent disease. The Journal of steroid biochemistry and molecular biology. 2002; 83(1-5):149–55.

40. Koehler KF, Helguero LA, Haldosén LA, Warner M, Gustafsson JA. Reflections on the discovery and significance of estrogen receptor β. Endocrine reviews. 2005; 26(3):465–78.

41. Laheri S, Ashary N, Bhatt P, Modi D. Oviductal glycoprotein 1 (OVGP1) is expressed by endometrial epithelium that regulates receptivity and trophoblast adhesion. Journal of assisted reproduction and genetics. 2018;35(8:1419–29.

42. Laheri S, Modi D, Bhatt P. Extra-oviductal expression of oviductal glycoprotein 1 in mouse: Detection in testis, epididymis and ovary. Journal of biosciences. 2017;42(1):69–80.

43. Langoi D, Pavone ME, Gurates B, Chai D, Fazleabas A, Bulun SE. Aromatase inhibitor treatment limits progression of peritoneal endometriosis in baboons. Fertility and sterility. 2013;99(3):656–62.

44. Li Y, Adur MK, Kannan A, Davila J, Zhao Y, Nowak RA, Bagchi MK, Bagchi IC, Li Q. Progesterone alleviates endometriosis via inhibition of uterine cell proliferation, inflammation and angiogenesis in an immunocompetent mouse model. PloS one. 2016; 11(10):e0165347.

45. Lin YJ, Lai MD, Lei HY, Wing LY. Neutrophils and macrophages promote angiogenesis in the early stage of endometriosis in a mouse model. Endocrinology. 2006;147(3):1278–86.

46. Livak KJ, Schmittgen TD. Analysis of relative gene expression data using real-time quantitative PCR and the 2− ΔΔCT method. Methods. 2001; 25(4):402–8.

47. McKinnon B, Mueller M, Montgomery G. Progesterone resistance in endometriosis: an acquired property?. Trends in Endocrinology & Metabolism. 2018.

48. Nisolle M, Casanas-Roux F, Wyns C, De Menten Y, Mathieu PE, Donnez J. Immunohistochemical analysis of estrogen and progesterone receptors in endometrium and peritoneal endometriosis: a new quantitative method. Fertility and sterility. 1994; 62(4):751–9.

49. Park JS, Lee JH, Kim M, Chang HJ, Hwang KJ, Chang KH. Endometrium from women with endometriosis shows increased proliferation activity. Fertility and sterility. 2009;92(4):1246–9.

50. Patel BG, Rudnicki M, Yu J, Shu Y, Taylor RN. Progesterone resistance in endometriosis: origins, consequences and interventions. ActaobstetriciaetgynecologicaScandinavica. 2017;96(6):623–32.

51. Pelch KE, Sharpe-Timms KL, Nagel SC. Mouse model of surgically-induced endometriosis by auto-transplantation of uterine tissue. Journal of visualized experiments: JoVE. 2012(59).

52. Pellegrini C, Gori I, Achtari C, Hornung D, Chardonnens E, Wunder D, Fiche M, Canny GO. The expression of estrogen receptors as well as GREB1, c-MYC, and cyclin D1, estrogen-regulated genes implicated in proliferation, is increased in peritoneal endometriosis. Fertility and sterility. 2012; 98(5):1200–8.

53. Pereira FE, Almeida PR, Dias BH, Vasconcelos PR, Guimarães SB, Medeiros FD. Development of a subcutaneous endometriosis rat model. Actacirurgicabrasileira. 2015 (1):6–12.

54. Plante BJ, Lessey BA, Taylor RN, Wang W, Bagchi MK, Yuan L, Scotchie J, Fritz MA, Young SL. G protein-coupled estrogen receptor (GPER) expression in normal and abnormal endometrium. Reproductive Sciences. 2012;19(7):684–93.

55. Qi QM, Guo SW, Liu XS. Estrogen Biosynthesis and Its Regulation in Endometriosis. Reproductive and Developmental Medicine. 2017; 1(1):55.

56. Qian H, Xuan J, Liu Y, Shi G. Function of G-protein-coupled estrogen receptor-1 in reproductive system tumors. Journal of immunology research. 2016;2016.

57. Rahimi S, Akaev I, Marani C, Chopra M, Yeoh CC. Immunohistochemical expression of different subtypes of cytokeratins by endometrial stromal sarcoma. Applied Immunohistochemistry and Molecular Morphology. 2018.

58. Rocha AL, Reis FM, Petraglia F. New trends for the medical treatment of endometriosis. Expert opinion on investigational drugs. 2012; 21(7):905–19.

59. Shao R, Cao S, Wang X, Feng Y, Billig H. The elusive and controversial roles of estrogen and progesterone receptors in human endometriosis. American journal of translational research. 2014; 6(2):104.

60. Simitsidellis I, Gibson DA, Saunders PT. Animal models of endometriosis: Replicating the aetiology and symptoms of the human disorder. Best Practice & Research Clinical Endocrinology & Metabolism.2018.

61. Somigliana E, Vigano P, Rossi G, Carinelli S, Vignali M, Panina-Bordignon P. Endometrial ability to implant in ectopic sites can be prevented by interleukin-12 in a murine model of endometriosis. Human Reproduction. 1999; 14(12):2944–50.

62. Taylor HS, Osteen KG, Bruner-Tran KL, Lockwood CJ, Krikun G, Sokalska A, Duleba AJ. Novel therapies targeting endometriosis. Reproductive sciences. 2011;18(9):814–23

63. Tirado-González I, Barrientos G, Tariverdian N, Arck PC, García MG, Klapp BF, Blois SM. Endometriosis research: animal models for the study of a complex disease. Journal of reproductive immunology. 2010;86(2):141–7.

64. Tosti C, Biscione A, Morgante G, Bifulco G, Luisi S, Petraglia F. Hormonal therapy for endometriosis: from molecular research to bedside. European Journal of Obstetrics & Gynecology and Reproductive Biology. 2017; 209:61–6.

65. Vigano P, Candiani M, Monno A, Giacomini E, Vercellini P, Somigliana E. Time to redefine endometriosis including its pro-fibrotic nature. Human Reproduction. 2017 Dec 1; 33(3):347–52.

66. Wetendorf M, DeMayo FJ. Progesterone receptor signaling in the initiation of pregnancy and preservation of a healthy uterus. The International journal of developmental biology. 2014;58:95.

67. Wingfield M, Macpherson A, Healy DL, Rogers PA. Cell proliferation is increased in the endometrium of women with endometriosis. Fertility and sterility. 1995;64(2):340–6.

68. Wu Y, Strawn E, Basir Z, Halverson G, Guo SW. Promoter hypermethylation of progesterone receptor isoform B (PR-B) in endometriosis. Epigenetics. 2006;1(2):106–11.

69. Xue Q, Lin Z, Cheng YH, Huang CC, Marsh E, Yin P, Milad MP, Confino E, Reierstad S, Innes J, Bulun SE. Promoter methylation regulates estrogen receptor 2 in human endometrium and endometriosis. Biology of reproduction. 2007; 77(4):681–7.

70. Yamanaka A, Kimura F, Takebayashi A, Kita N, Takahashi K, Murakami T. Primate model research for endometriosis. The Tohoku journal of experimental medicine. 2012; 226(2):95–9.

71. Zanatta A, Pereira RM, Rocha AM, Cogliati B, Baracat EC, Taylor HS, Motta EL, Serafini PC. The relationship among HOXA10, estrogen receptor α, progesterone receptor, and progesterone receptor B proteins in rectosigmoid endometriosis: a tissue microarray study. Reproductive Sciences. 2015 (1):31–7.

72. Zhao Y, Li Q, Katzenellenbogen BS, Lau LF, Taylor RN, Bagchi IC, Bagchi MK. Estrogen-induced CCN1 is critical for establishment of endometriosis-like lesions in mice. Molecular Endocrinology. 2014; 28(12):1934–47.

